# Relationship between constitutive and acute gene regulation, and physiological and behavioral responses, mediated by the neuropeptide PACAP

**DOI:** 10.1101/2021.03.29.437579

**Authors:** Dana Bakalar, Sean Sweat, Gunner Drossel, Sunny Z. Jiang, Babru S. Samal, Nikolas Stroth, Wenqin Xu, Limei Zhang, Haiying Zhang, Lee E. Eiden

**Affiliations:** Section on Molecular Neuroscience, National Institute of Mental Heath - Intramural Research Program, Bethesda, MD. NIH, USA; Department of Physiology, Autonomous National University of Mexico (UNAM) Medical School, Mexico City, Mexico

## Abstract

Since the advent of gene knockout technology in 1987, insight into the role(s) of neuropeptides in centrally- and peripherally-mediated physiological regulation has been gleaned by examining altered physiological functioning in mammals, predominantly mice, after genetic editing to produce animals deficient in neuropeptides or their cognate G-protein coupled receptors (GPCRs). These results have complemented experiments involving infusion of neuropeptide agonists or antagonists systemically or into specific brain regions. Effects of gene loss are often interpreted as indicating that the peptide and its receptor(s) are required for the physiological or behavioral responses elicited in wild-type mice at the time of experimental examination. These interpretations presume that peptide/peptide receptor gene deletion affects only the expression of the peptide/receptor itself, and therefore impacts physiological events only at the time at which the experiment is conducted. A way to support ‘real-time’ interpretations of neuropeptide gene knock-out is to demonstrate that the wild-type transcriptome, except for the deliberately deleted gene(s), in tissues of interest, is preserved in the knock-out mouse. Here, we show that there is a cohort of genes (constitutively PACAP-Regulated Genes, or cPRGs) whose basal expression is affected by constitutive knock-out of the *Adcyap1* gene in C57Bl6/N mice, and additional genes whose expression in response to physiological challenge, in adults, is altered or impaired in the absence of PACAP expression (acutely PACAP-Regulated Genes, or aPRGs). Distinguishing constitutive and acute transcriptomic effects of neuropeptide deficiency on physiological function and behavior in mice reveals alternative mechanisms of action, and changing functions of neuropeptides, throughout the lifespan.

## Introduction

Neuropeptides were discovered, characterized chemically, and their receptors profiled pharmacologically over the course of the decades from 1910-1970 [1]. cDNAs for transcripts encoding the neuropeptides and their G-protein coupled receptors were obtained during the 1980s and 90s. The last known neuropeptide receptors to be cloned were the opiate receptors MOR, DOR and KOR, with other peptide-liganded GPCRs subsequently de-orphanized [2, 3]. The technology for gene knockout was developed by the Capecchi lab in 1987 [4], and originally applied to investigation of homeobox genes and their influence on mammalian development, including that of the central nervous system. Knock-out technology, applied to individual peptide and peptide receptor genes throughout the 1990’s and early 00’s (e.g., [5]) allowed a more definitive causality to be applied to the roles and functions of neuropeptides/neuropeptide receptors than was possible pharmacologically with chemical agonists and antagonists.

However, as Hokfelt and colleagues point out, the era of molecular biology did not lead automatically to the clear identification of physiological roles for individual peptides [6]. This may be partly because, as peptides are released under conditions of high neuronal activity (e.g., stress), their main contribution to neuroendocrine regulation is to allow ‘efficacy in extremis’, rather than primary action in basal physiology [7]. A second reason is that neuropeptides may act not only immediately and directly, following peptide secretion/release via synaptic and paracrine interactions with their receptors [8], but also indirectly, by influencing the chemoanatomy of brain, peripheral nervous system, and endocrine tissues during development. Interpreting the results of loss-of-function experiments following genetic editing of neuropeptide and neuropeptide receptor genes may therefore require accounting for both immediate (direct) effects of secreted neuropeptides and downstream activation of their receptors, and for indirect effects mediated through altered expression of mRNAs affecting the expression and function of other proteins. Thus, loss of function in neuropeptide-ablated mice could be attributed either to the absence of the peptide itself, at time of experiment, or to developmental or other indirect effects of neuropeptide loss or absence. Useful interpretation of knock-out experiments in adult mice, especially in a translational context, clearly requires assessment of the time of impact of neuropeptide deficiency. Distinguishing effects of neuropeptide deficiency on the basal wild-type transcriptome, which might indirectly affect physiological and behavioral response patterns, from deficiencies in real-time gene regulation required for execution of physiological and behavioral responses, is an important step in that assessment.

An early example of perturbation of brain function secondary to the immediate loss of peptide expression and release was provided by the Young laboratory studying the effects of knock-out of oxytocin (Oxt) and vasopressin (AVP) receptor genes in the mouse [9]. Constitutive knock-out of either the AVP type 1 or Oxt receptors did not affect freezing behavior following fear conditioning. However, forebrain-specific knock-out of the oxytocin receptor after weaning resulted in reduction in Avpr1 expression in the central nucleus of the amygdala, and subsequent impairment in fear conditioning. The authors interpreted their results as indicating a compensation for loss of Oxt or AVP receptors preserving patent fear conditioning responses if occurring early enough in development, but a subsequent lack of compensation in young, post-weaning mice, during a period in which Oxt receptor is needed for maintenance of AVP receptor gene transcription presumably required for patent function of circuits involved in fear learning. Given the intimate association between Oxt and AVP function in learning and memory circuits, and in the development of such circuitry in mammals, it is perhaps unsurprising that abolishing expression of Oxt, AVP, or the receptors for either, might have unequally penetrant effects at different onsets of peptide/receptor deficiency in the mouse.

Similar concerns about compensatory effects of PACAP and VIP mediated through the actions of their shared receptors PAC1, VPAC1 and VPAC2 prompted May and colleagues to examine the expression of PACAP, VIP, PAC1, VPAC1 and VPAC2 mRNA as a function of constitutive (from inception) knock-out of either PACAP or VIP [10]. In this case, deficiency in expression of either PACAP or VIP did not affect the expression of any of the four remaining components of this overlapping and potentially self-compensating system in the brain. However, these experiments beg the question of whether neuropeptide constitutive knock-out has long-term indirect effects on the transcriptome, or immediate effects on either cellular excitability or cell plasticity. To investigate the mechanism(s) of PACAP action in the mouse we examined, in wild-type (WT) and PACAP knockout (PACAP KO) mice, the transcriptomes of several brain areas in which PACAP is released from nerve terminals, and the adrenal gland, at which PACAP is the sole identified neuropeptide neurotransmitter.

Based upon the results described here, we suggest that interpretation of the physiological role(s) of neuropeptides in the diffuse neuroendocrine system, including the brain, based on phenotypic presentation in constitutive-knockout mice, requires prior examination of the entire transcriptome of such animals, at least in tissues relevant to the behaviors or physiological functions studied. Furthermore, developmental and acute effects of neuropeptide deficiency on the brain transcriptome can be correlated with altered phenotypic outcomes, both physiological and behavioral. This can contribute to understanding the dual functional roles of central nervous system neuropeptides in nervous system development and in real time neurotransmission in adult mice.

## Materials and Methods

### Transgenic and knockout mice

PACAP- and PAC1-deficient C57BL6/N mice were generated as previously described [11-13]. Wild type (WT) refers to corresponding C57BL6/N strains at time of experimentation. PACAP^fl/fl^ mice were generated as follows: The CRISPR-Cas9 system with a donor plasmid was used to generate the floxed PACAP (*Adcyap1*) allele on a C57BL/6N background. CRISPR-Cas9 mediated the specific double-strand DNA cleavage on both 5’ and 3’ of the coding sequence (CDS) of the PACAP gene where two LoxP sites are located in the edited gene within ‘floxed’ mice. The donor plasmid contained a LoxP site at both a 5’ and 3’ site within the CDS of *Adcyap1* (P38), located in exon 5 of the *Adcyap1* gene [14] to permit PACAP excision as in the constitutive knockout [11]. LoxP sites were flanked by *Adcyap1* sequences mediating homologous recombination of the LoxP-CDS (*Adcyap1*)-LoxP region into the *Adcyap1* gene of the transgenic mouse. sgRNAs were designed using MIT Guide RNA design software (Feng Zhang laboratory, previously available at crispr.mit.edu) and CCTop (https://cctop.cos.uni-heidelberg.de), a CRISPR/Cas9 target online predictor were used to minimize off-target recombination and optimize specificity for the flanking sequences of the CDS of PACAP. The 5’sgRNA was designed to avoid the predicted splicing signal and corresponds to the sequence segment (5’-agtcacagtattcccgccagCGG-3’) located in the intron upstream of Exon 5. The 3’sgRNA was designed to target the sequences (5’-CCCactggttgcaggggcaattc-3’) in exon 5 of the *Adcyap1* gene to facilitate the insertion of LoxP 567 base pairs after the STOP codon following Cas9 cleavage of the double-stranded genomic DNA. Capital letters in the targeting sequences indicate the PAM sites. A donor plasmid was generated on a TA vector backbone and contains an ∼1.3 kb sequence homologous to the *Adcyap1* gene upstream of the 5’LoxP and a 1.5 kb sequence homologous to the *Adcyap1* gene downstream of the 3’LoxP site.

sgRNAs were purchased from IDT as Alt-R crRNA and tracrRNA oligos, which were mixed in dH_2_O at a ratio of 1:2 and annealed to generate sgRNA in a thermocycler (95°C for 5 min and then ramp down to 25°C at a rate of 5°C /min). gRNA and Cas9 protein (IDT) were mixed at room temperature for 15 minutes to form a ribonucleoprotein complex with subsequent addition of donor plasmid DNA to generate the injection mix. Each component had a final concentration of 20 ng/µl in 1xTAE buffer. The injection mix was then passed through a Millipore filter (UFC30VV25). The final injection solution was injected into zygotes to produce founder mice in the NIMH-IRP Transgenic Core Facility. Accurate CRISPR-Cas9 cleavage of the target site and insertion of the LoxP sites was confirmed by PCR amplification, and sequencing, of the region.

Camk2α CRE PACAP^fl/fl^ mice are a cross between mice expressing CRE under the Camk2α promoter and mice with LoxP sites flanking exon 4 of the *Adcyap1* gene. These animals were bred as described in Jiang, 2017 [15]. Vgat CRE PACAP^fl/fl^ mice are a cross between the PACAP floxed mice described above and a commercially available Vgat-ires-CRE line (B6J.129S6(FVB)-*Slc32a1*^*tm2(cre)Lowl*^/MwarJ, Jackson Labs)

Mice used for experiments described here were between eight and 32 weeks of age. Both male and female mice were used.

### Dissection of brain and peripheral tissue for qRT-PCR and microarray experiments

At least three male and three female mice of each genotype, aged 10-20 weeks, were used for microarray analysis and qRT-PCR assay. Mice were sacrificed by cervical dislocation and whole brains were quickly removed from the skull and rinsed in cold 1× PBS buffer. Two lateral cuts were made, at the optic chiasm and above the cerebellum and brainstem respectively, to obtain a coronal tissue block. Hypothalamus was microdissected along the anterior commissure from one hemisphere to the other using a blade and fine-tipped forceps. Whole hippocampus was removed after peeling off the cerebral cortex from the same coronal tissue block. The prefrontal cortex was dissected according to The Mouse Brain in Stereotaxic Coordinates, Paxinos and Franklin, 2001 [16] (Bregma 1.5 to 0.5 mm) and cerebellar cortex was also collected from each hemisphere. Both adrenal glands, and a portion of the median lobe of the liver were collected from the same animals. All dissected tissue was quickly frozen in Dry Ice and stored at -80 C until RNA extraction.

### RNA extraction for qRT-PCR and microarray

Total RNA was extracted using an miRNeasy Mini kit (QIAGEN, Cat#: 217004, Valencia, CA, USA), according to the manufacturer’s instructions. Briefly, frozen tissue (20 mg or less) from brain, adrenal glands, or liver was disrupted using a plastic pestle in 700μl lysis buffer per sample, then centrifuged at 12,000 × g for 15 min at 4° C. Following centrifugation, the supernatant was transferred to a new microcentrifuge tube, and 1.5 volumes of 100% ethanol was added and mixed by pipetting. The mixed sample was transferred to an RNeasy mini-spin column and centrifuged for 30 s at 8,000 × g. The RNA bound to the column membrane was washed out with RWT buffer then with RPE buffer. The RNA was then dissolved in 30-40μl of diethylpyrocarbonate (DEPC)-treated water. RNA concentration was evaluated by absorbance at 260 nm using the NanoVue Plus spectrophotometer (Biochrom US, Holliston, MA, USA). RNA purity and integrity were further determined by Bioanalyzer, and only RNA with RNA Integrity Number (RIN) greater than 8 was used in microarray or qRT-PCR experiments. RNA was stored at -80 °C until use.

### cDNA synthesis and qRT-PCR

Single-stranded cDNAs were synthesized with the Superscript III first strand cDNA synthesis kit (Cat#: 18080051, Thermo Fisher Scientific, Waltham, MA, USA), according to the kit protocol, using 1-2 mg total RNA. qRT-PCR protocols were as reported previously [17, 18]. All Taqman gene expression assay kits were purchased from Thermo Fisher Scientific (Waltham, MA, USA). Taqman probes were designed using Partek Genomic Suite software, also used for the microarray analysis, to align probe sequences with microarray results. Detailed information for each probe is listed in Table 1.

**Table 1:** Taqman gene expression assays for qRT-PCR.

| Gene symbol | Assay ID |
| --- | --- |
| *Adcyap1* | Mm00437433_m1 |
| *Pttg1* | Mm00479224_m1 |
| *Xaf1* | Mm01245815_m1 |
| *Mid1* | Mm04933165_m1 |
| *Pla2g4e* | Mm00625711_m1 |
| *Gapdh* | Mm99999915_g1 |

The qRT-PCR reactions were performed in a BioRad iCycler as follows: 95° C hold for 20 s followed by 40 cycles of 95° C, denaturation for 3 s, and 60° C annealing and extension for 30 s. qRT-PCR analyses for each mRNA transcript were performed using the 2^-ΔΔCt^ method [19]. In the present study, data are presented as the fold-change in expression of each transcript, normalized first to the internal *Gapdh* transcript and then relative to the reference tissue within each tissue type. The cycle threshold (Ct) was defined as the number of cycles required for the fluorescent signal to cross the threshold. ΔCt was determined as [mean of the duplicate Ct values for each gene] – [mean of the duplicate Ct values for *Gapdh*]. ΔΔCt represented the difference between the paired tissue samples (e.g., PACAP KO vs. WT), as calculated by the formula ΔΔCt = [ΔCt for each transcript in PKO tissue - ΔCt for each transcript in reference WT tissue]. The fold change for each transcript in specific tissue was expressed as 2^-ΔΔCt^ [17-19].

### Microarray analysis

Analysis was conducted in the NHGRI-NINDS-NIMH Microarray Core (Abdel Elkahloun, Director) using its established protocols (available at https://ma.nhgri.nih.gov/) and Clariom-S mouse microarray chips (Affymetrix).

### Bioinformatics

Microarray data was analyzed using Partek Genomics Suite. CEL files from five groups of microarrays (two from hypothalamus, and one each from adrenal gland, cerebellum, and hippocampus) were imported separately for each experiment, using RMA normalization. Within each experiment, differentially expressed genes were determined by ANOVA, with pairwise contrasts between groups of interest. To compare hypothalamic transcriptomes of constitutive PACAP KO and WT mice, we designated genes as differentially expressed with a fold-change of plus-or-minus 2 and a raw p-value of p <= 0.01. The relatively relaxed raw p-value was chosen rather than an adjusted p-value to avoid type II error. Analysis of replication across multiple data sets provided additional power. Other thresholds used are identified and described in text and/or figure legends. Where raw expression values are shown, data are given as Log2 expression value, normalized in Partek via RMA normalization of CEL files from all microarray analysis experiments.

### RNAscope in-situ hybridization

In-situ hybridization was conducted using the RNAScope Multiplex Fluorescent V2 Assay (ACD Bio), as directed in the manual. Briefly, fresh-frozen mouse brains were sectioned sagittally on a cryostat to 16µM. Sections were mounted on Superfrost Plus slides (Fisher Scientific) and dried on slides at -20 degrees. Slides were fixed with 4% paraformaldehyde for 15 minutes at 4 degrees, then dehydrated in an ethanol gradient. Hydrogen peroxide and protease treatments, probe hybridization and signal development proceeded as described in the manual. Probes were Mm-Pttg1-C3 (Cat No. 560291-C3), and Mm-Xaf1(Cat No. 504181). These were developed with Opal 620 and Opal 690 fluorophores (Akoya Biosciences). Imaging was performed in the NIMH Systems Neuroscience Imaging Resource (SNIR) on a Stellaris confocal microscope.

## Results

### Hypothalamic transcriptome of PACAP-deficient compared to wild-type mice

Differences in the hypothalamic transcriptomes of WT and PACAP-deficient mice were assessed. In the first experiment, designated “Hypothalamus Experiment 1”, RNA was harvested from hypothalamus of four PACAP KO male mice and three WT male mice. Nineteen genes differed between PACAP KO and WT males at a fold-change of +/- 2, p <= 0.01. This includes 4 upregulated and 16 downregulated genes, including *Adcyap1*. Given the sexually dimorphic nature of the hypothalamus, we repeated this experiment with an equal number of male and female mice (n = 3 male and n = 3 female animals of each genotype, “Hypothalamus Experiment 2”). We identified transcripts whose expression was altered regardless of sex by performing sex by genotype ANOVA and excluding from consideration genes with significant interactions at p < 0.01. In Hypothalamus Experiment 2, 16 transcripts were altered (5 upregulated, 11 downregulated) in PACAP KO versus WT animals.

As shown in Figure 1, a subset of 13 transcripts, including *Adcyap1* itself, was altered in PACAP KO versus WT mice in both hypothalamic experiments. We will refer to these as **cPRGs** (**c**onstitutively **P**ACAP-**R**egulated **G**enes). Ten transcripts (*Pttg1, Mid1, 4833420G17RIK, Pisd-ps3, Cd59a, Gm5148, Cyb5r3, Tmem260, and Cetn4*, as well as *Adcyap1* itself) showed significantly lower expression in PACAP-deficient mice and three transcripts (*Pla2g4e, Mrpl20*, and *Xaf1*) showed significantly higher expression in constitutively PACAP-deficient mice. Although none of the identified cPRGs are regulated differently in mice of different sexes, we did find several sex-dependently expressed transcripts in hypothalamus, in good agreement with the literature [20] (Figure 1 Supplemental).

**Figure 1.**
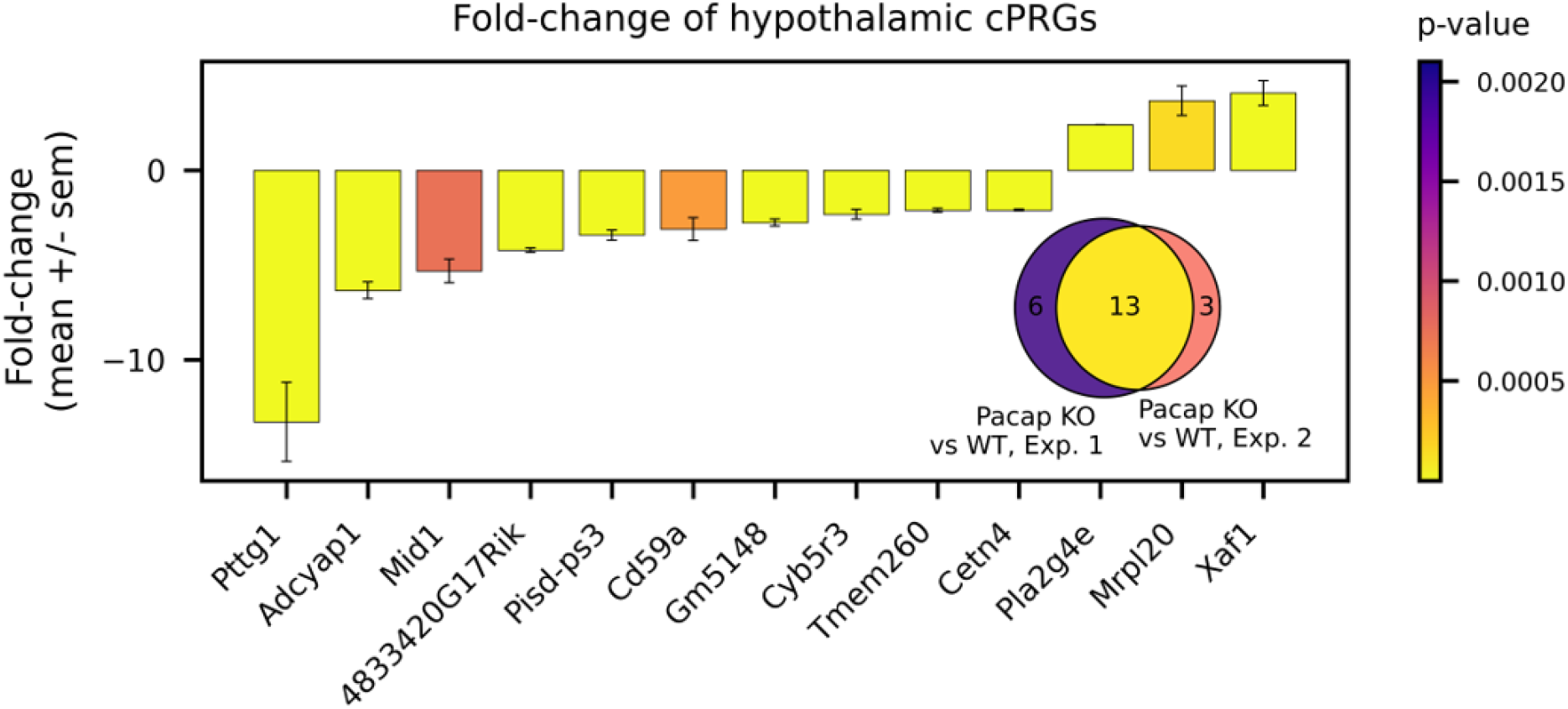
Transcripts differentially expressed in hypothalamus of constitutively PACAP-deficient mice. Microarray analysis reveals ten genes which are differentially expressed (p < 0.01, fold-change +/- 2 or greater) between non-stressed PACAP KO and non-stressed WT mice in both hypothalamic microarrays. Hypothalamus experiment 1: N = 3 male wildtype and N = 3 male PACAP KO. Hypothalamus experiment 2, N = 3 males and N = 3 females of each genotype. Bar graph shows the mean fold-change (+/- standard error of the mean) of PACAP KO hypothalamus versus WT hypothalamus across both experiments. Color bar indicates p-value of comparison. Inset Venn depicts overlap between the two experiments.

Identification of constitutively PACAP-regulated transcripts was extended to the cerebellum and hippocampus, which express modest amounts of *Adcyap1* in excitatory (hippocampus) or inhibitory (cerebellum) neurons [21]. Because of these lower baseline levels of *Adcyap1*, the threshold for fold-change was shifted to +/- 1.1 in these tissues (Figure 2A). The loss of power resulting from lowered threshold was compensated for by identifying genes which replicate, that is, which are regulated in multiple experiments. In hippocampus, 63 genes were differentially regulated in PACAP KO versus WT animals. In cerebellum, 58 genes met the criteria.

**Figure 2.**
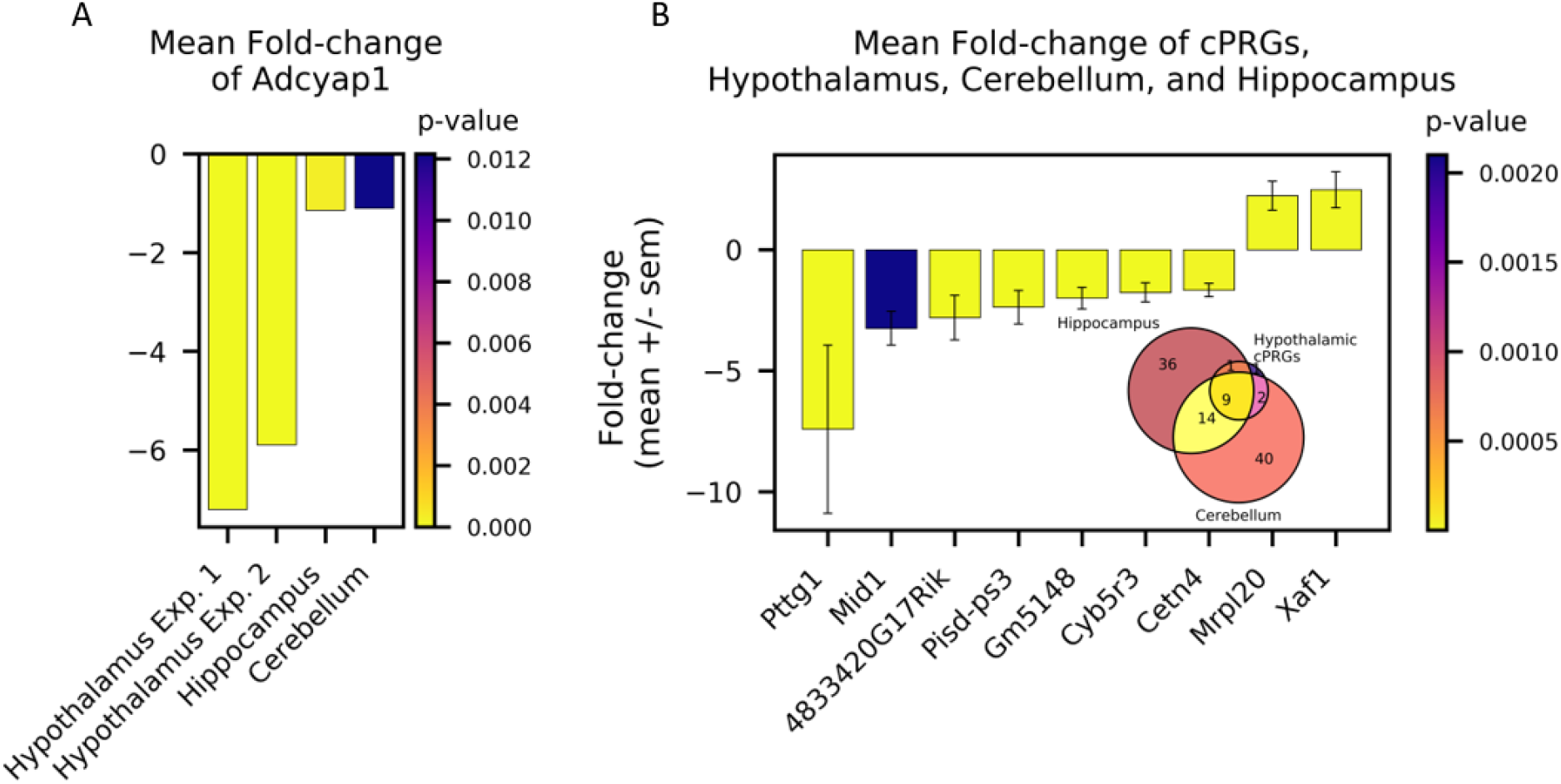
Constitutively PACAP-regulated genes across three brain regions. **A)** Fold-change in *Adcyap1* expression in PACAP-deficient mice in various tissues. Bar graph shows fold-change of PACAP KO mice versus WT mice within each experiment. Color bar indicates p-value of comparison. Fold-change in *Adcyap1* expression varies across tissues/brain regions despite identical deletion of the *Adcyap1* gene at all sites due to differences in the intrinsic expression of the *Adcyap1* transcript and the threshold for transcript detection by microarray analysis (see text). **B)** Constitutively PACAP-regulated genes (cPRGs) identified by microarray analysis in hippocampus and cerebellum (N = 3 males and N = 3 females of each genotype in each tissue) and hypothalamus, as described previously. Genes were identified as cPRGs if they were differentially expressed between PACAP KO and WT mice (p < 0.01, fold-change +/- 1.1 or greater) in PACAP knockout versus WT cerebellum and hippocampus and differentially expressed p < 0.01, fold-change +/- 2 or greater in both hypothalamic experiments. Color bar indicates p-value of comparison.

A subset of 10 of the differentially expressed transcripts in both hippocampus and cerebellum overlapped with the hypothalamically identified cPRGs, without being differentially expressed in male and female tissue (Figure 2B: *4833420G17Rik, Cetn4, Cyb5r3, Gm5148, Irgm2, Mid1, Pisd-ps3, Pttg1, Pyurf*, and *Xaf1*). A further fourteen genes (*1700048O20Rik, Abdh1, Adat2, Gvin1, Irgm2, Itgb3bp, Kbtbd11, Lacc1, Nmrk1, Pyurf, Slc25a37, Slc39a2, Stxbp2, Zfp125*) were commonly regulated between hippocampus and cerebellum but were not hypothalamic cPRGs. The cPRG transcripts most dramatically down-regulated (2-fold or more) are *Pttg1*, a transcriptional activator associated with development [22]; *Mid1*, implicated in structural brain defects, Huntington’s and Alzheimer’s pathology [23], *4833420G17Rik*, a predicted protein (orf) of unknown function, and *Pisd-ps3*, phosphatidylserine decarboxylase pseudogene 3, regulated in aging [24]. *Mrpl20* and *Xaf1* are genes encoding known proteins of unknown function in the nervous system.

*Adcyap1* itself does not register as significantly downregulated in hippocampus (Figure 2A). This is an artefact of the threshold for detection of rare transcripts by microarray. As *Adcyap1* is expressed at already very low levels in hippocampus and cerebellum, compared to hypothalamus (Figure 3A), its apparent fold-change is constrained by the dynamic range of microarray in these tissues (Figure 3A, B). Indeed, qRT-PCR for expression of *Adcyap1* in hypothalamus, cerebellum, and hippocampus in WT vs. PACAP KO mice shows a main effect of genotype (F(1,53), p < 0.0001, with significant differences between PACAP KO and WT mice in hypothalamus (p = .0026), cerebellum (p = 0.014), and hippocampus (p = .0081) (Figure 3B), but not in adrenal gland (not shown), in which *Adcyap1* is not detectable in our assay (average Ct value of 31.9 in WT and 32.8 in KO mice).

**Figure 3.**
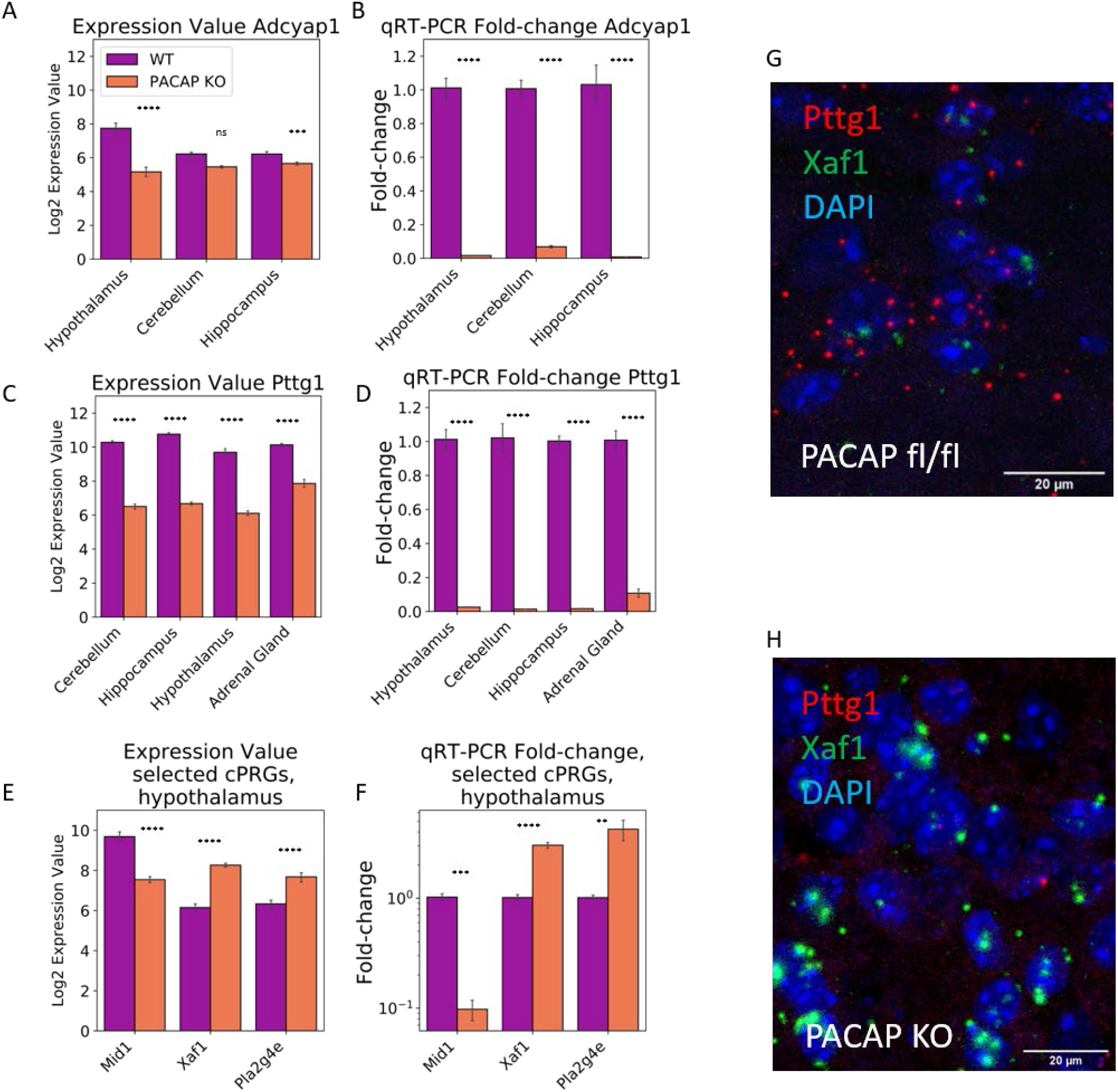
Comparison of expression values for transcripts encoding the cPRGs *Pttg1, Mid1, Xaf1, Pla2g4e*, and *Adcyap1*, and independent verification of regulation assessed by qRT-PCR. **A)** Log2 expression values of *Adcyap1* in non-stressed PACAP knockout versus wildtype mice of both sexes, in all microarray experiments. Expression of *Adcyap1* is higher in wildtype hypothalamus than in other neural tissues. Comparison of wildtype and PACAP KO mice within each tissue shows that tissues with lower baseline *Adcyap1* expression have smaller reductions in *Adcyap1* expression in PACAP KO versus wildtype mice (Stars indicate significance of ANOVA comparing WT expression values to KO expression values. In hypothalamus, p = 3.39e-13, fold-change = -6.55; in cerebellum p = 3.17e-4, fold-change = -1.14; but not in hippocampus p = 1.22e-2, fold-change -1.09). **B)** qRT-PCR, with larger dynamic range, reveals significant decreases in *Adcyap1* expression in all PACAP producing tissues. Two-way ANOVA revealed a significant main effect of genotype F(1,33) = 476.9, p < 0.0001. Post-hoc Šídák’s comparisons shows differences between PACAP KO and wildtype animals in hippocampus, hypothalamus, and cerebellum, p < 0.0001 in all tissues. Bar graph depicts mean and standard error of fold-change calculated using the 2^−ΔΔCt^ technique with *Gapdh* as control. **C)** Log2 expression values of *Pttg1* in non-stressed PACAP knockout and wildtype mice of both sexes, in all microarray experiments. Putative cPRG *Pttg1* is highly expressed in all tissues studied and is significantly reduced (ANOVA, p < 0.0001) in PACAP KO versus wildtype animals in all tissues. **D)** qRT-PCR confirms significant loss of *Pttg1* in hypothalamus, cerebellum, hippocampus, and adrenal gland (F(1,43) = 967.5, p < 0.0001, Post-hoc Šídák’s show p < 0.0001 for all comparisons). Bar graph depicts mean and standard error of fold-change calculated using the 2^−ΔΔCt^ technique with *Gapdh* as control. **E)** Log2 expression values, in hypothalamus, of *Mid1, Xaf1*, and *Pla2g4e*, identified as cPRGs, in hypothalamus. ANOVA indicates significant differences (p < 0.0001) in each gene in hypothalamus. **F)** qRT-PCR confirms that transcriptomic alterations are significant in the direction found in microarrays. Multiple t-tests with Bonferroni correction shows significant differences between WT and KO for *Mid1* (t(13) = 10.97, adjusted p < 0.0001), *Xaf1* (t(13) = 8.89, P = 0.000002) and *Pla2g4e* (t(13) = 3.156, p = 0.022). Bar graph depicts mean and standard error of fold-change calculated using the 2^−ΔΔCt^ technique with *Gapdh* as control. **G, H)** RNAscope in-situ hybridization visualizing mRNA for *Pttg1* (red) and *Xaf1* (green) are depicted in lateral hypothalamus of WT (G) and PACAP KO (H) mouse. The decrease in *Pttg1* expression and increase in *Xaf1* expression are visually evident in the PACAP KO mouse.

The most consistently and dramatically downregulated cPRG, from the microarray experiments described above, was *Pttg1*. Its microarray expression value is significantly depressed in PACAP KO versus WT animals in all tissues studied (Figure 3C). We confirmed its decreased expression in hippocampus, hypothalamus, and cerebellum, and adrenal gland of PACAP KO mice using qRT-PCR. Two-way tissue x genotype ANOVA revealed a significant effect of genotype (F(1,43) = 967.5, p < 0.0001), with Šídák’s multiple comparisons showing differences at p < 0.0001 in all tissues (Figure 3D).

To confirm additional cPRGs (microarray expression in hypothalamus in Figure 3E), we used qRT-PCR to assess expression of *Mid1, Xaf1*, and *Pla2g4e* in hypothalamus. Unpaired t-tests with Benjamini, Krieger, and Yekutieli False Discovery Rate compensation for multiple testing were performed for each gene in each tissue, comparing PACAP KO and WT mice. All comparisons were significant (hypothalamic results shown in Figure 3F, remaining results in Table 2).

**Table 2:** qRT-PCR analysis of cPRGs.

| Gene | Tissue | P value | t ratio | df | q value |
| --- | --- | --- | --- | --- | --- |
| *Mid1* | Hypothalamus | <0.000001 | 10.97 | 13 | <0.000001 |
| *Xaf1* | Hypothalamus | <0.000001 | 8.892 | 13 | 0.000002 |
| *Pla2g4e* | Hypothalamus | 0.003913 | 3.561 | 12 | 0.004517 |
| *Xaf1* | Hippocampus | <0.000001 | 8.892 | 13 | 0.000002 |
| *Mid1* | Hippocampus | 0.0001 | 6.212 | 10 | 0.000118 |
| *Xaf1* | Cerebellum | 0.000021 | 7.502 | 10 | 0.000036 |
| *Mid1* | Cerebellum | 0.000082 | 6.367 | 10 | 0.000116 |

### Effects of restraint stress on the hypothalamic transcriptome in wild-type and PACAP-deficient mice

Examination of transcripts up-regulated by restraint stress, versus the non-stressed condition in WT and in PACAP KO mice allows detection of transcripts whose stress-associated induction is PACAP-dependent. In both PACAP KO and WT mice, a number of transcripts, including the immediate-early genes (IEGs) *Nr4a1, Nr4a3, Sgk1, Cyr61*, and *Nfkbia* were upregulated significantly (p < 0.01, fold-change +/- 1.5) by one hour of restraint stress (Figure 4A), demonstrating that there is an intact stress transcriptomic response in PACAP-deficient mice.

**Figure 4.**
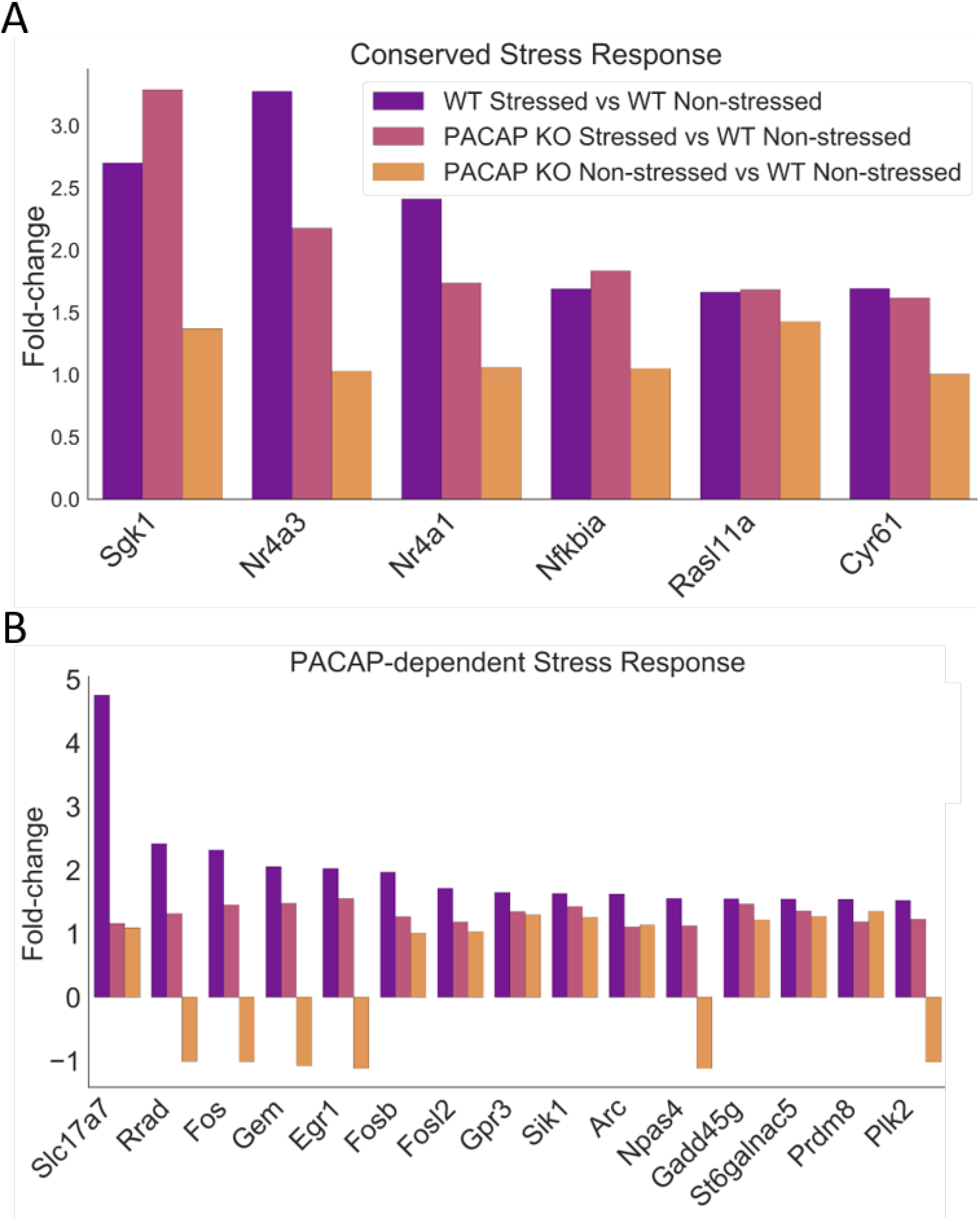
Effects of stress on hypothalamic transcriptome of PACAP KO versus WT mice. **A)** Conserved stress response genes are regulated by stress in WT animals (comparison of WT stressed to WT non-stressed, purple, p < 0.01) and in PACAP KO mice (comparison of PACAP KO stressed to wildtype non-stressed, pink, p < 0.01). These genes do not differ between WT and PACAP KO mice without stress (yellow). **B)** A PACAP-dependent stress response is indicated by genes which are upregulated in response to stress among wildtype mice (comparison of wildtype stressed to wildtype non-stressed, p < 0.01, fold-change +/- 3 or greater shown) but fail to be recruited in PACAP KO mice. These genes do not differ between WT and PACAP KO mice without stress (yellow).

Several other IEGs, however, were PACAP-dependent, as they were upregulated by stress in wild-type animals and not in PACAP knockout mice (Figure 4B). These were *Rrad, Fos, Gem, Egr1, Fosb, Fosl2, Arc, Npas4, Gadd45g*, and *Plk2. Fos* and *Egr1* have previously been shown to be PACAP-dependent upon one hour of restraint stress in hypothalamus and adrenal gland [25]. Because these genes are not altered in the baseline transcriptome of the PACAP KO mouse (Figure 1), lack of stress-induced regulation is due not to regulation of these target genes during development, but to acute effects of loss of PACAP transmitter function during stress in hypothalamus, making these genes **a**cutely **P**ACAP-**R**egulated **G**enes (aPRGs).

### Effects of PACAP knockout on adrenal gland transcriptome

The adrenal gland is a neuroendocrine organ in which *Adcyap1* mRNA is expressed at very low levels, but which expresses both specific (PAC1) as well as relatively non-specific (BAM22P) receptors for PACAP. It is copiously innervated by the splanchnic nerve, which allows PACAP release onto adrenomedullary chromaffin cells throughout the lifespan, beginning in early postnatal life, and may therefore be altered by the loss of PACAP signaling throughout development [26-28]. We identified cPRGs in adrenal gland, in which PACAP KO and WT mice differed significantly (p < 0.01, fold-change +/- 1.5). Of the 93 differentially regulated genes (70 downregulated and 23 upregulated), 7 genes had been identified as cPRGs in our hypothalamic analysis. These genes are *4833420G17Rik, Cd59a, Cyb5r3, Mrpl20, Pisd-ps3, Pttg1*, and *Tmem260*. Three genes were members of the catecholamine biosynthesis pathway (*Th, Pmnt*, and *Hand1*). These final three may not be cPRGs, strictly speaking, as developmental and adult functions at the adrenomedullary synapse may significantly overlap.

The adrenal glands are the major peripheral effectors of hypothalamic stress signaling [29, 30], and PACAP is a known regulator of these circuits [25, 31, 32]. We therefore examined the stress-regulated genes in adrenal gland that were either PACAP-dependent (expression increased significantly, p < 0.01, at a fold-change of 1.5 or more in WT but not PACAP KO mice) or conserved between PACAP KO and WT animals (increased in both). Figure 5A shows a *subset* of the 192 genes (those most upregulated by 3 hours of restraint stress, fold change > 3) in both PACAP-deficient and WT adrenal gland. Figure 5B depicts a *subset* of the 303 PACAP-dependent adrenal transcripts, for which WT animals showed increased transcription (fold-change > 3) but PACAP KO animals did not, and for which expression did not differ between non-stressed PACAP KO and WT mice. As in hypothalamus, genes regulated acutely by PACAP (aPRGs) can be distinguished from those whose expression is altered by constitutive loss of PACAP (cPRGs): Only two genes were altered in the non-stressed comparison and in the stressed comparison (*Scgn* and *Pcsk1*).

**Figure 5.**
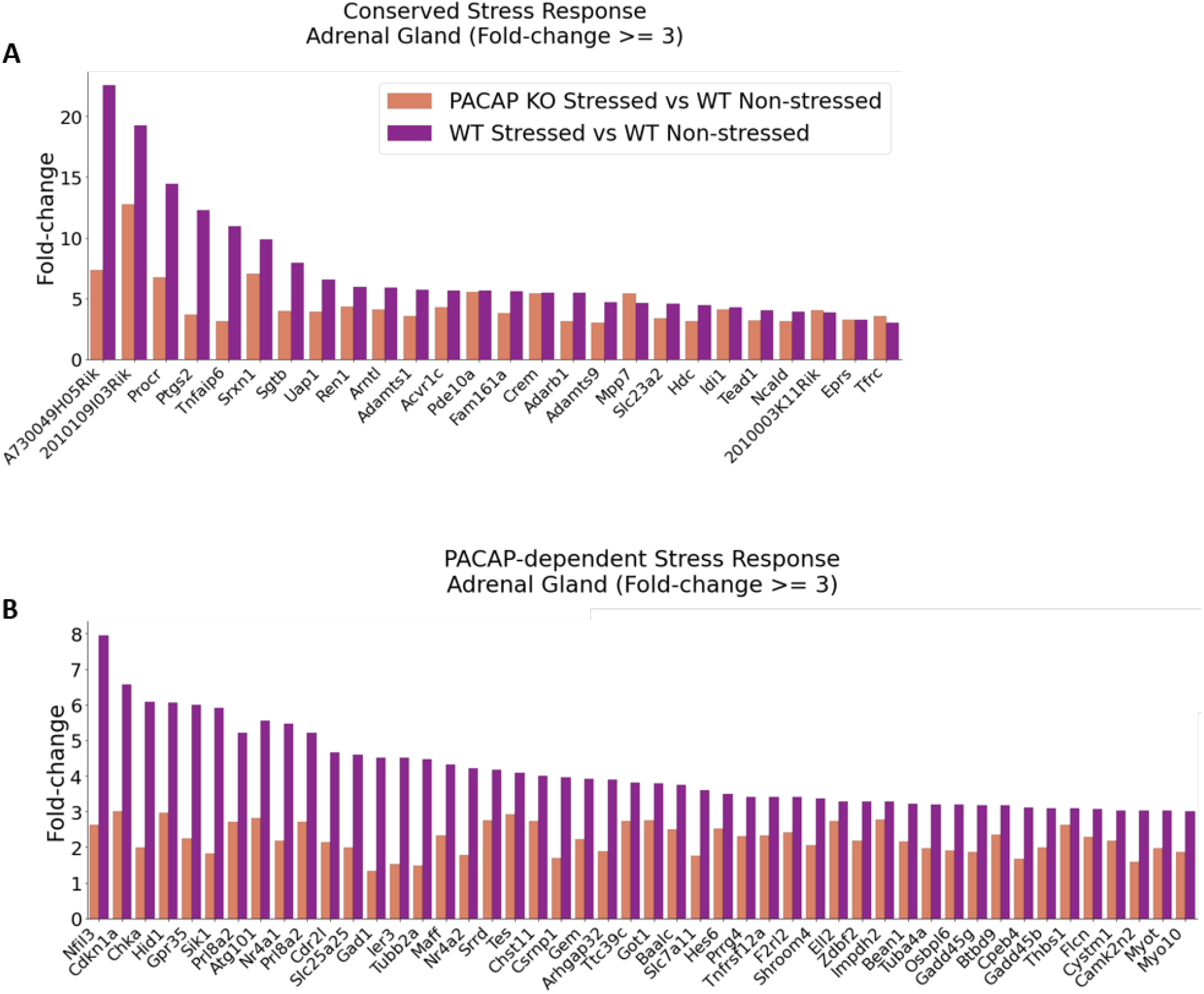
Effects of PACAP knockout and stress on adrenal transcriptome. Conserved stress response genes are regulated by stress in WT animals (comparison of WT stressed to WT non-stressed, p < 0.01, fold-change +/- 3 or greater) and in PACAP KO mice (comparison of PACAP KO stressed to wildtype non-stressed, p < 0.01, fold-change +/- 3 or greater). **B)** A PACAP-dependent stress response is indicated by genes which are upregulated in response to stress among WT mice (comparison of WT stressed to WT non-stressed, p < 0.01, fold-change +/- 3 or greater) but fail to be recruited in PACAP KO mice.

One gene regulated by stress in WT but not PACAP KO mice is immediate early gene 3 (*Ier3*). *Ier3* has also been referred to as PACAP-regulated gene 1, or *Prg1*, because it was identified as a gene up-regulated by PACAP in rodent pancreatic tumor cells [33]. Another regulated gene is *Stc1*: PACAP regulates expression of both *Ier3* and *Stc1* in bovine chromaffin cells in culture [34-36] and in cultured rodent cortical neurons [37]. We confirmed these aPRGs with qRT-PCR (Figure 6A’ and A’’). The alteration in *Stc1* is adrenal-specific and does not occur in hypothalamus in either microarray (Figure 6B) or qRT-PCR analysis (Figure 6B’, B’’).

**Figure 6.**
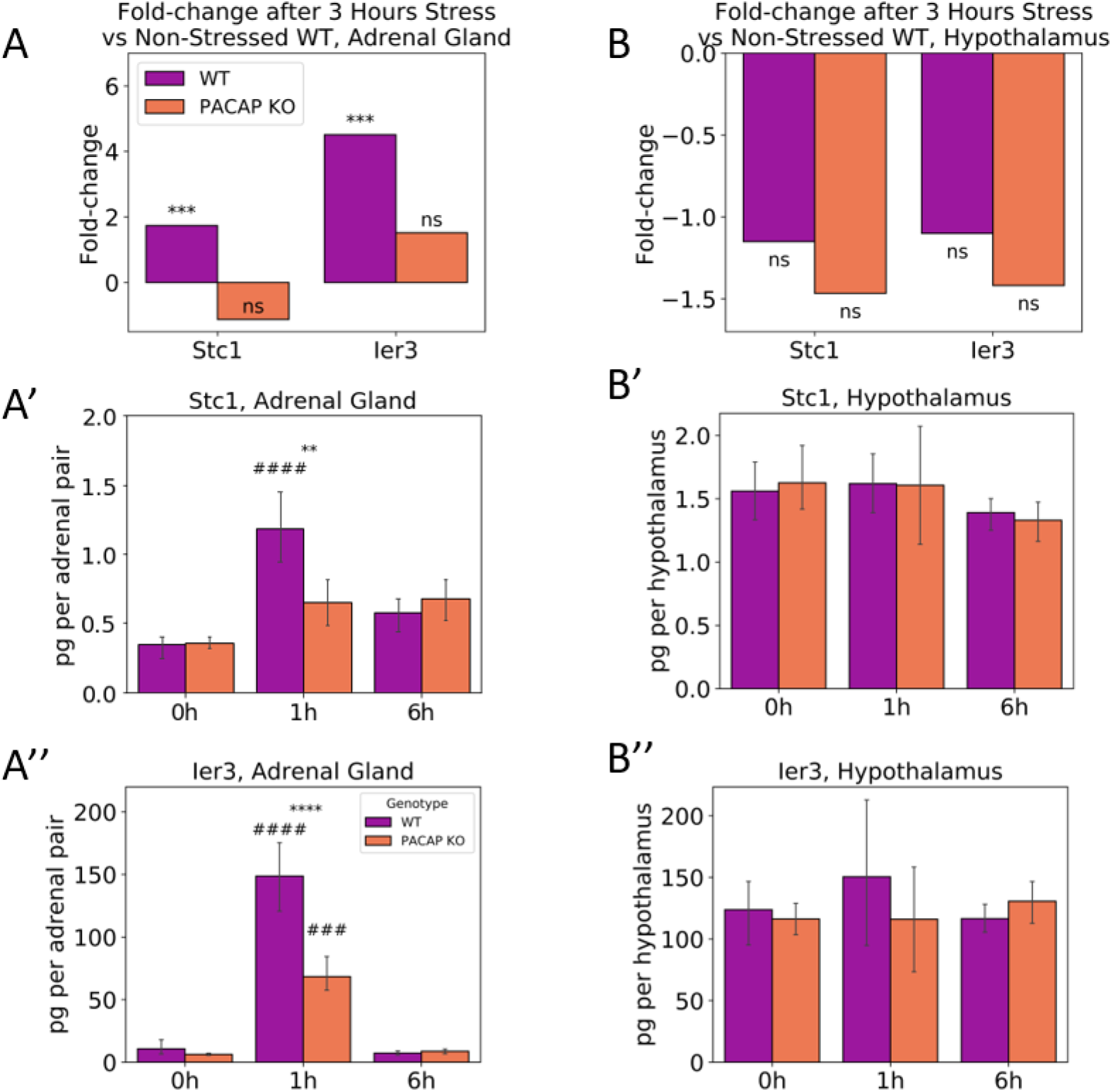
Adrenal gland, stress, and regulation of the aPRGs Stc1 and Ier3. **A)** Log2 expression values of IEGs *Ier3* and *Stc1* in adrenal gland microarray show increased *Ier3* and *Stc1* following 3 hours of restraint in wildtype but not PACAP knockout mice, relative to non-stressed wildtype mice. **A’)** Two-way ANOVA analysis of qRT-PCR for adrenal *Stc1* shows a significant interaction between time restrained and genotype F(2,25) = 6.99, p = 0.0039: While there was no difference between non-stressed wildtype and PACAP KO mice, Šídák’s test showed significant increases in WT (p < 0.0001), but not in PACAP KO (p = 0.419) adrenal gland following 1 hour restraint stress. Wildtype animals, however, had significantly higher *Stc1* expression (p = 0.0035) than PACAP KO mice at 1-hour stress. At 6 hours of stress, *Stc1* levels had dropped relative to 1-hour stress significantly in wildtype (p = 0.0015) mice, returning to baseline levels. Asterisks indicate significant difference between genotypes within timepoints, while pound signs indicate significant difference between untreated and 1 hour stressed animals within each genotype. **A’’)** qRT-PCR shows that immediate early gene *Ier3* is increased after 1-hour restraint stress in wildtype and PACAP KO adrenal gland. Two-way ANOVA shows a significant interaction between time restrained and genotype F(2,25) = 15.60, p < 0.0001: While there was no difference between non-stressed wildtype and PACAP KO mice, Šídák’s test showed significant increases in both WT, p < 0.0001, and PACAP KO p = 0.0002 adrenal gland after 1 hour stress, compared to non-stressed animals. Wildtype animals, however, had significantly higher *Ier3* expression after stress (p < 0.0001) than PACAP KO mice. By 6 hours of restraint, *Ier3* levels had dropped relative to 1-hour stress significantly in wildtype (p < 0.001) and PACAP KO (p = 0.0004) animals, returning to baseline levels. Asterisks indicate significant difference between genotypes within timepoints, while pound signs indicate significant difference between untreated and 1 hour stressed animals within each genotype. **B)** Log2 expression values of IEGs *Ier3* and *Stc1* in hypothalamus show no significant difference in wildtype and PACAP KO regulation of *Ier3* or *Stc1* following 3 hours of restraint relative to non-stressed wildtype mice. **B’, B’’)** qRT PCR confirms no alteration of *Stc1* or *Ier3* in either wildtype or PACAP KO mice after 1 or 6 hours of stress.

### Behavioral and physiological responses in PACAP-deficient mice

Finally, we examined the possible links between the transcriptomic impact of PACAP deficiency on cPRGs and aPRGs, and mouse spontaneous and stress-elicited behaviors. Figures 7A and 7B depict the acute effects of PACAP deficiency on corticosterone levels, and on weight loss secondary to decreased food intake (hypophagia), as measured 24 hours after three hours of restraint stress. As reported previously for chronic stress effects on feeding [38, 39], PACAP deficiency abrogates weight loss in both male and female PACAP-deficient mice, and attenuates cortisol elevation after three hours of restraint. In contrast, repetitive jumping appears to be a constitutively established behavior not triggered by a specific physiological event such as restraint stress: With no discrete triggers, PACAP KO mice perform stereotyped repetitive jumping [40] throughout the dark cycle, up to 147 times per half hour period in the current study, while control mice (PACAP^fl/fl^) repetitively jumped significantly less (Figure 7C, 7C’).

**Figure 7.**
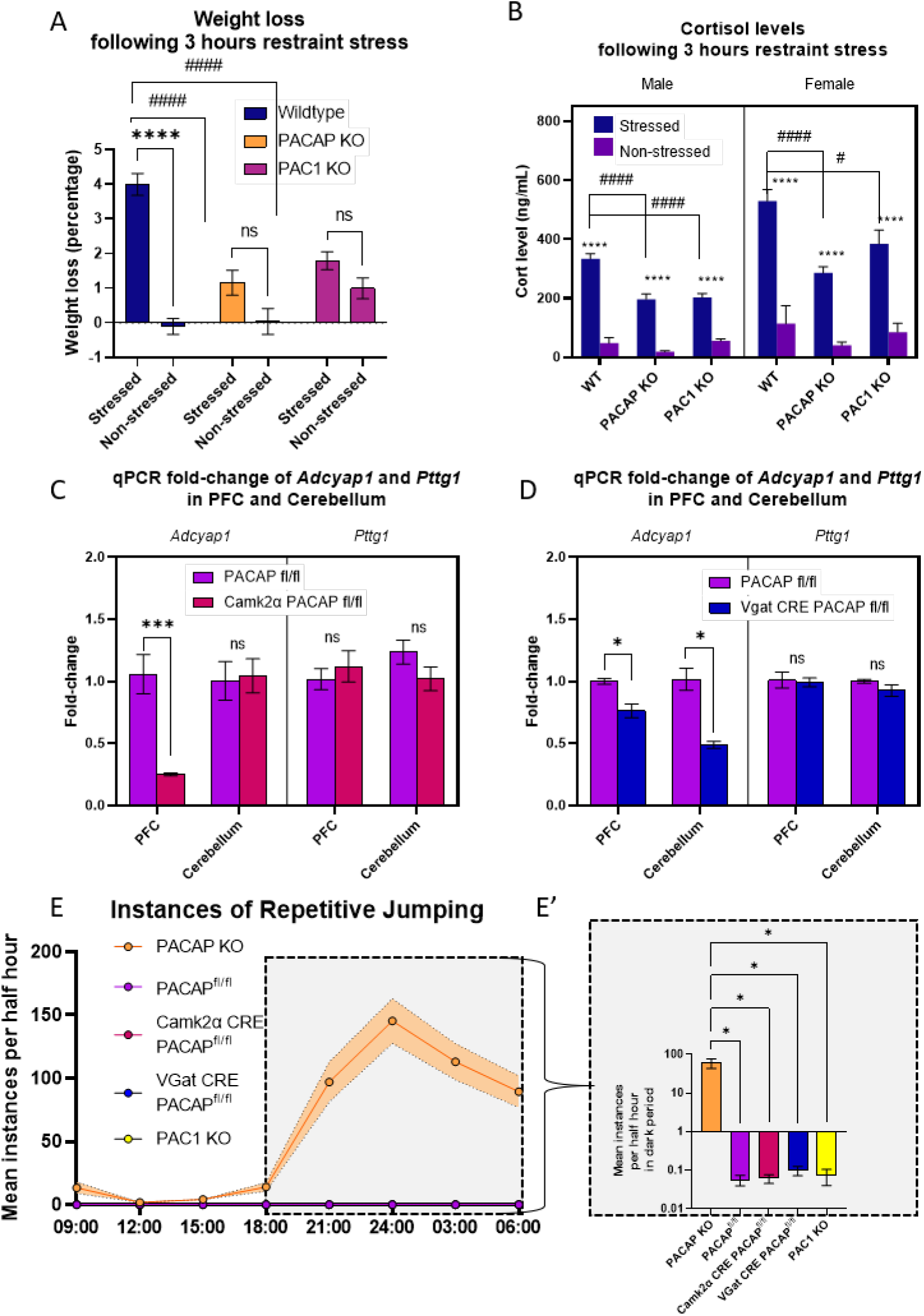
Physiological and Behavioral phenotypes: Hypophagia, Cortisol, Repetitive jumping. **A)** Blunting of stress-induced hypophagia occurs in both male and female PACAP KO mice. Three-way ANOVA found no main effect of sex and no interactions between sex and either stress or genotype, so sexes were combined and analyzed with 2-way ANOVA. There were main effects of Stress (F(2,94) = 53.76, p < 0.0001), of Genotype (F(2,94) = 7.21, p = 0.0012) and a significant Stress by Genotype interaction (F(2,94) = 14.6, p < 0.0001). Šídák’s multiple comparisons test showed that while WT mice lost significant weight following stress (p < 0.0001), neither PACAP nor PAC1 KO mice did. Therefore, stressed WT mice lost significantly more weight than stressed PACAP KO or PAC1 KO mice (p < 0.0001 for both). **B)** Cortisol increases following 3 hours of restraint stress in PACAP KO, PAC1 KO, and WT mice of both sexes (3-way ANOVA, Main effect of Stress F (1, 88) = 397.2, p < 0.0001). Tukey’s multiple comparisons test showed significant increases after stress in all groups (p < 0.0001, stars). However, the increase is blunted in both male and female PACAP KO mice (hashes; Male WT Stressed vs Male PACAP KO Stressed, p < 0.0001; Male WT Stressed vs Male PAC1 KO stressed, p < 0.0001; Female WT Stressed vs Female PACAP KO Stressed, p < 0.0001; Female WT Stressed vs Female PAC1 KO stressed, p = 0.012). **C)** Lack of jumping is correlated with non-depletion of the cPRG *Pttg1*. In prefrontal cortex of Camk2α CRE PACAP^fl/fl^ mice, compared with PACAP^fl/fl^ control mice, *Adcyap1* was significantly reduced (2-way ANOVA, Šídák’s multiple comparisons test p = 0.0003), but *Pttg1* was not. In cerebellum, neither gene was altered in Camk2α CRE PACAP^fl/fl^ mice relative to controls. **D)** In Vgat CRE PACAP^fl/fl^ mice, *Adcyap1* was significantly reduced in both PFC (2-way ANOVA, Šídák’s multiple comparisons test p = 0.015) and cerebellum (2-way ANOVA, Šídák’s multiple comparisons test p <0.0001), but Pttg1 was not altered relative to controls in either tissue. **E, E’)** While constitutive PACAP knockout mice perform a repetitive jumping behavior at high rates (yellow line), none of the other tested animals (PACAP^fl/fl^ controls, PAC1 KO, Camk2α CRE PACAP^fl/fl^, and Vgat CRE PACAP^fl/fl^) jump. Repetitive jumping over the course of 24 hours is shown in 7E, while average number of repetitive jumping bouts per half hour during the dark cycle is shown in E’. Brown-Forsythe ANOVA test revealed a significant effect of genotype (F(4,11) = 12.96), with Dunnett’s T3 multiple comparisons test showing significantly higher jumping in PACAP KO than all other groups (p = 0.036 for all comparisons).

### Behavioral phenocopying in conditional PACAP- and in constitutive PAC1-deficient mice

Because acute effects of PACAP transmission occur primarily via the PAC1 receptor, we examined the phenocopying of the above features of PACAP KO mice by constitutive PAC1-deficient mice (PAC1 KO). The abrogation of stress-induced weight loss is phenocopied in PAC1-deficient mice, suggesting that the effect depends on intact PACAP-PAC1 signaling. Attenuation of cortisol increase was also evident in the PAC1 KO mouse. On the other hand, repetitive jumping, does not occur in PAC1-deficient mice (Figure 7E, F), suggesting that repetitive jumping stems from long-term effects of PACAP deficiency arising from altered expression of cPRGs.

We wished to further correlate the potential role(s) of cPRGs and aPRGs in PACAP control of physiological responses and behavioral phenotypes in the mouse. Anticipating that cPRGs may have an indirect effect on function due to changes in the basal transcriptome, we sought to understand how early in development PACAP control of the transcription of genes such as *Pttg1*, and of behavioral outputs like repetitive jumping might ensue. We created two conditional knockout mice to investigate, by crossing a newly-developed conditional PACAP knock-out mouse (PACAP^fl/fl^), to two CRE-driver mice; the Camk2α PACAP^fl/fl^ mouse in which CRE expression, and therefore PACAP knock-out, occurs under the control of the Camk2α promoter, which becomes active in forebrain during early postnatal life [41], and the Vgat CRE PACAP^fl/fl^ mouse, in which PACAP-knockout ensues postnatally in cerebellum [42]. This allows us to examine the behavioral and transcriptomic effects of PACAP knockout in postnatal excitatory and inhibitory neurons.

We measured, using qRT-PCR, levels of *Adcyap1* and *Pttg1* in prefrontal cortex (PFC) and cerebellum of each line. In Vgat CRE PACAP^fl/fl^ animals, *Adcyap1* was reduced in both tissues, although more dramatically in cerebellum where more *Vgat* expression is expected. In Camk2α PACAP^fl/fl^ mice, *Adcyap1* transcript levels are reduced by more than eighty percent in prefrontal cortex, and not at all in cerebellum, where PACAP expression is largely limited to expression in inhibitory neurons (Purkinje cells) [21, 43]. *Pttg1* expression was unaffected by conditional PACAP-deficiency in either tissue in either knockout line (Figure 7C, D). It may be that control of *Pttg1* (and other cPRG) transcription by PACAP is supported by PACAP release from cell types in which these promoters are not active. Nevertheless, the complete lack of effect on *Pttg1* expression in brain regions in which post-natal loss of PACAP expression is profound, does suggest that PACAP plays an earlier-than-postnatal role in regulation of aPRGs such as *Pttg1*.

Behaviorally, while constitutive PACAP KO mice perform hundreds of repetitive jumping bouts per half-hour period, neither control mice (PACAP floxed, CRE negative) nor either conditional knockout have the repetitive jumping phenotype (Figure 7E, E’). This suggests that *Pttg1*, or other aPRGs, may be functionally important in this phenotype, although reverse genetic analysis will be required to demonstrate this.

## Discussion

The neuropeptide PACAP, like several other neuropeptides, is expressed during embryonic development prior to complete development of the nervous system [44-46], and throughout adult life in both peripheral and central nervous systems. Neuropeptides are expressed at locations indicating neurotransmitter function upon co-release with acetylcholine from preganglionic sympathetic and parasympathetic neurons, with glutamate and other neuropeptides from sensory neurons, and with either glutamate or GABA from both projection and interneurons of the CNS [14, 21, 47-50]. How neuropeptides such as PACAP act as instructive or trophic factors, at various stages of development, is less well understood [45, 51-53]. Here, we have compared the transcriptomes of wild-type mice and constitutively PACAP-deficient mice (i.e. lacking PACAP since inception) in central and peripheral nervous system, and in both the basal state, and after undergoing response to restraint stress. Transcriptomic analysis after both constitutive and conditional PACAP knock-out reveals two distinct patterns of gene expression influenced by PACAP, with separate linkages to physiological response to stress and to spontaneous behavior.

A core group of transcripts were identified that are up- or down-regulated in PACAP KO compared to WT mice in all brain regions examined, including hypothalamus, hippocampus and cerebellum, under basal conditions. These genes, prominently including *Pttg1* and *Xaf1*, are *not* regulated in tissues/brain regions in which PACAP knockout occurs late in development (in Camk2α CRE PACAP^fl/fl^ or Vgat CRE PACAP^fl/fl^ mice). These are referred to as constitutive PACAP-regulated genes, or cPRGs, indicating that their regulation by PACAP action not as a neurotransmitter, but as a neurotrophic factor during development. Regulation of cPRGs, rather than aPRGs (see below) is correlated with repetitive jumping in PACAP-deficient mice: The selective loss of PACAP from CNS in either excitatory or inhibitory neurons, occurring later in development in VGAT CRE PACAP^fl/fl^ or Camk2α CRE PACAP^fl/fl^ mice, is accompanied neither by regulation of cPRGs, nor by repetitive jumping.

It is of interest that repetitive jumping is also not phenocopied in PAC1-deficient mice. Other receptors for PACAP exist—VPAC1 and VPAC2--which have been implicated in PACAP action in vivo, although not during development [54]. MRG3B, the BAM22P receptor, which acts as a low-affinity receptor for multiple neuropeptides including PACAP [55], is a less-characterized but plausible candidate as a receptor through which PACAP might exert its developmental actions. Although this receptor is restricted to primary sensory neurons in the adult [56], it may be more widespread earlier in development. It would be illuminating to test whether various cell culture correlates of PACAP action during development (e.g. [57-59] are likewise phenocopied in neurons cultured from PAC1 ko mice.

Acute PACAP-regulated genes, or aPRGs, are identified here as transcripts which are induced during a physiological event, in this case restraint stress, in WT but not in PACAP KO mice. These genes are equally expressed in both WT and PACAP KO mice prior to stress exposure. In the periphery, chromaffin cells of the adrenal medulla are targeted by cholinergic/PACAPergic innervation, and PACAP has a critical =role in mediating stress transduction by the chromaffin cell [11, 54, 60, 61]. We have previously examined the PACAP-regulated transcriptome in both primary chromaffin cells, and in pheochromocytoma cells induced to differentiate by this neuropeptide [36, 62, 63]. Genes regulated immediately upon exposure to PACAP (i.e. what we refer to here as aPRGs) include *Ier3* and the neuroprotective calcium storage modulator stanniocalcin (*Stc1*) [64]. *Ier3* (also known as PRG-1, is the first gene reported to be PACAP-regulated, in the pancreatic AR4-2J acinar pancreatic cell line [65]. It is worth noting that fundamental developmentally-determined alteration of the nervous and endocrine systems, via aPRGs, could also lead to hypofunction in the stress response, despite the lack of obvious alteration in adrenal structure and function in the basal state [66].

Restraint stress causes up-regulation of multiple genes in the hypothalamus, as in the adrenal medulla. Unlike cPRGs, aPRGS appear to be highly tissue-specific upon comparison of the hypothalamic and adrenal transcriptomes after restraint stress. We have previously reported on deficits in endocrine and behavioral stress responding in PACAP-deficient C57Bl6/N mice, and we and others have generally interpreted the effects of PACAP deficiency as contributory to phenotype primarily due to its absence at the time of stress responding [25, 38, 39, 61, 67-69]. However, only rescue experiments or highly temporally restricted PACAP knockout, can establish whether or not the effects of PACAP deficiency in adult animals are due to its action solely at the time of experimentation in the adult, or neuronal impairment secondary to the differential expression of one or more cPRGs.

The whole-mouse knockout of the PACAP gene has yielded significant information about the roles of this ‘master regulator’ of the stress response (see [60] and references therein). Complementary experiments in conditional knock-out mice in which PACAP is eliminated in specific brain regions at or near the time of experimentation will reveal more about the dual role of this neuropeptide in trophic/developmental specification of neuronal and endocrine properties on the one hand, and real-time neurotransmitter effects in specific physiological responses on the other. These further investigations may also illuminate mechanisms of dysregulation of stress responding throughout the lifespan.

The present results suggest two discrete roles for PACAP in regulation of nervous system function. One of these is ‘real-time’, presumably neurotransmitter-mediated, control of the mouse stress response, and is closely associated with acute PACAP-regulated genes (aPRGs). This occurs both centrally and peripherally, as reflected in the PACAP-dependent transcriptomic response to stress in both hypothalamus and adrenal gland [70, 71]. A distinct, second role for PACAP is manifested during development, and is associated with the PACAP-dependent expression of constitutively expressed PACAP-regulated genes (cPRGs). At least one effect of early developmental transcriptomic control by PACAP appears to involve motor output, as evidenced by the development of repetitive jumping behavior following early developmental PACAP deficiency, which does not occur when knockout is induced later in development.

The mechanisms of each of the two discrete types of PACAP regulation, and the circuits in which PACAP signaling is critical for each, remain to be explored. Further experiments in conditional PACAP- and PAC1-deficient mice, and precise correlation of time of onset of PACAP deficiency and emergence of the functional effects of down-regulation of PACAP target genes, will be helpful in this regard. The absence or presence of PACAP can affect cerebellar development via modulation of mitogenesis and cortical migration of cells destined for the inner granular layer in early postnatal life [72-75]. However, these effects are apparently transient, as obvious cerebellar architectural defects have not been reported in the adult PACAP-deficient mouse [52]. Likewise, PACAP signaling can affect cortical neuronal progenitor cell cycle dynamics ex vivo [45, 76], yet obvious effects of PACAP deficiency on adult neocortical cytoarchitecture have not been reported. We have previously hypothesized that these effects, while not required for normal brain development, represent latent PACAP signaling capabilities that may be important in neuroprotection during prenatal or neonatal hypoxia or toxic insult such as occurs in fetal alcohol exposure [66]. In addition, early loss of PACAP expression has important effects on metabolism and thermal regulation during early post-natal development, although whether these effects are phenocopied by PAC1 deficiency has not been established [77]. In contrast, PACAPergic signaling in adult mammals appears to be consequential for mediating stress responding, and to occur via discrete real-time post-synaptic effects, likely mediated through both transcriptional and non-transcriptional mechanisms ([78, 79] and references therein).

The distinction between two modes of PACAP signaling to the genome, reported here, are relevant to both developmental and adult actions of PACAP. It remains to be determined if the two discrete types of neuropeptide function, constitutive and acute, exist for other neuropeptides expressed in the mammalian brain during both development and in the adult, and, if so, whether diverse signaling mechanisms, and even different sets of neuropeptide-responsive receptors, are required for each, as appears to be the case for PACAP control of cPRG and aPRG transcripts across the lifespan and in different regions of the central and peripheral nervous systems.

## Acknowledgements

We gratefully acknowledge NIMH transgenic core facility for generation of PACAP^fl/fl^ mice. Thanks to Michelle Sung for her help with genotyping PACAP^fl/fl^ mice. This work was supported by the National Institute of Mental Health Intramural Research Program, Project 1ZIAMH002386 to L.E.E.

## Figures

**Supplemental Figure 1.**
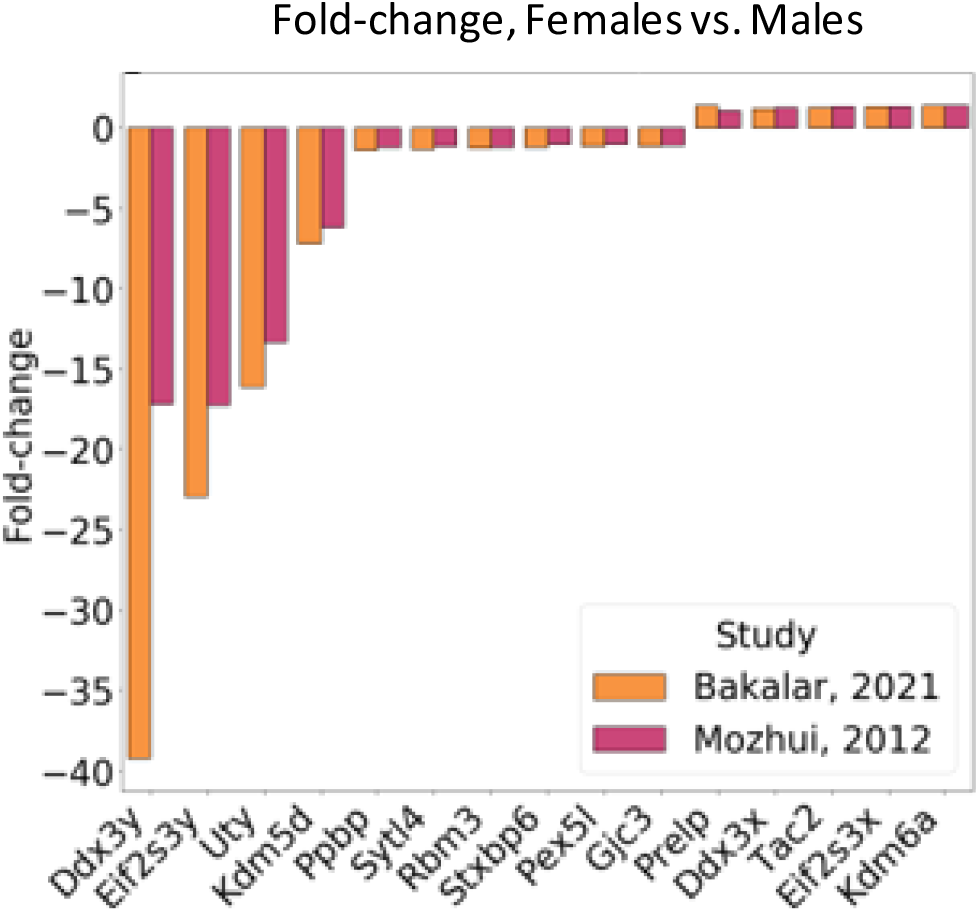
Identification of sex-dependent PACAP-regulated genes in the hypothalamus. Comparison of expression of male and female hypothalamic transcriptomes derived in this study to literature-reported sex-dependent gene expression in mouse brain. Following a two-way sex by genotype ANOVA, genes in our hypothalamic experiment 2, including male and female animals, were selected if females and males differed significantly (p < 0.01) in expression value. Data from Mozhui, 2012 [20], comparing hypothalamic transcripts of 39 pairs of adult BDX-strain mice, was retrieved and genes with a significant difference in expression between males and females (p < 0.01) were extracted. Fifteen genes were differentially regulated by sex in both experiments. These genes matched well in both direction and magnitude between the studies.

